# Longitudinal analysis of vaccine-associated alterations in the faecal microbiota of layer chickens using a vaccination schedule representative of commercial practice

**DOI:** 10.64898/2026.06.30.735456

**Authors:** Anum Ali Ahmad, Kris Hogan, Laura Glendinning

## Abstract

The gut microbiota is crucial for immune development and overall health in chickens. In commercial production, birds routinely receive multiple vaccines during early life. While individual vaccines are known to affect microbial composition, the impact of complex, multi-vaccine programs, as used in the poultry industry, is not well understood. This pilot longitudinal study examined the impact of multiple live and inactivated vaccines, given at commercially relevant times from an early age, on gut microbial diversity and composition in layer chickens. We characterised microbiota profiles using 16S rRNA gene sequencing at pre- and post-vaccination timepoints across different vaccine groups. Overall, microbial diversity remained stable across most vaccines, indicating strong resilience of the gut microbiota to repeated immunological interventions. Differential abundance analyses didn’t identify significant changes in taxa following vaccination. The gut microbial community also showed stability after the completion of the vaccination schedule. These preliminary findings underscore the robustness of the chicken gut ecosystem and lay a foundation for future research into microbiome-vaccine interactions and their implications for poultry health, immunity, and production efficiency.

## Background

The poultry industry is the leading source of animal protein for human consumption. In 2025, global poultry meat production exceeded 151 million metric tons, and egg production reached 99 million metric tons (FAO 2025). This rapid expansion has led to the development of high-density poultry farms, which increases the risk of disease outbreaks. Vaccination is crucial in poultry health management, as it protects against a range of diseases that threaten bird welfare and productivity.

The immune system and gut microbiota develop an interdependent relationship from birth, establishing long-term immune homeostasis. In vertebrates, about 70-80% of immune cells reside in the gut, interacting with the microbiome to maintain gut integrity, support immune maturation, and influence overall health (Wiertsema *et al*. 2021). Similarly, increasing evidence in chickens indicates that vaccines can modulate microbial composition through immune activation, while the microbiota can affect vaccine responsiveness by shaping mucosal and systemic immunity (Beck, Zhao, and Erf 2024, Liu PY *et al*. 2024). However, in modern poultry houses, young chicks lack transmission of maternal microbiota and primarily acquire microbiota from the environment, which potentially influences their responses to vaccines (Shterzer *et al*. 2023). Understanding the impact of vaccines on gut microbiota is essential for improving chicken health and vaccine efficacy.

Most research has focused on the impact of single vaccines on gut microbial communities of chickens, usually under experimental or controlled settings (Das *et al*. 2021, Khan *et al*. 2024, 2025). These studies indicate that vaccination can alter microbial diversity, shift bacterial populations, and temporarily disrupt gut balance. However, they only offer a limited perspective on vaccine-microbiome interactions. In commercial production systems, layer chicks routinely receive multiple live and inactivated vaccines at various developmental stages, from early life to sexual maturity. This regime protects birds against diverse pathogens throughout their production cycle. Yet, the cumulative effects of these complex vaccination schedules on gut microbial development are not well understood.

Understanding these interactions is crucial for layer chickens, where long-term productivity relies on strong immune function, efficient nutrient utilisation, and stable gut ecology. Disruptions in early-life microbiota have been linked to poor growth, reduced immune competence, and higher susceptibility to gut disorders (Simon *et al*. 2016). Moreover, commercial rearing conditions, characterised by environmental changes, stress, and repeated immunological challenges, may influence microbiome responses to vaccination (Akhtar and Zaman 2026). Despite the widespread use of multi-vaccine programs, no longitudinal studies to date have systematically evaluated how the administration of different vaccines in commercial settings impacts the gut microbiota dynamics in layer chickens.

Therefore, this pilot longitudinal study aimed to explore the influence of multiple live and inactivated vaccines on gut microbiota in layer chickens by a vaccination schedule representative of commercial practice. This work provides an important insight into microbiome resilience, host-microbe-vaccine interactions, and broader implications for poultry health and performance.

## Methodology

### Study design and sample collection

Faecal samples were collected from Hy-Line GP-11 chickens maintained for breeding purposes at the National Avian Research Facility, The Roslin Institute. The chickens were undergoing routine vaccinations as they transitioned into breeding and egg-production flocks, consistent with standard commercial vaccine practices. The birds were maintained under PUN039 (Breeding and Maintenance of Hy-Line Brown), housed in accordance with the Animals (Scientific Procedures) Act 1986 (ASPA) Code of Practice and subject to Animal Welfare and Ethical Review Body (AWERB) subcommittee approval.

The chicks were housed in four pens and wing-tagged for individual identification. All birds received a standard commercial starter crumb/mash from weeks 0 to 6, followed by rearer pellets from weeks 7 to 14. All birds were fed *ad libitum*. Lighting and temperature were managed in accordance with the recommendations provided in the Hy-Line Brown Commercial Layers Management Guide. The number of birds per pen varied throughout the trial (weeks 4-7: n=47 birds/pen; weeks 8-14: n=30 birds/pen) due to the removal of birds as they grew. However, the birds included in the study remained in their originally assigned pens throughout the trial period.

Three birds per pen were randomly selected for longitudinal sampling, and the same birds were followed throughout the study. Individual birds served as subsamples, and the pens served as the true independent biological replicates (n = 4). Vaccines were administered through drinking water, which provided an oral route of exposure. The birds were vaccinated in accordance with standard commercial practices, using the timing and vaccine combinations commonly used for layer breeders. Vaccinations were administered at weeks 4, 7, 9, and 12 according to the manufacturer’s recommended dosages. At week 4 (W4), three vaccines, Nobilis ND Clone 30, Nobilis Gumboro D78, and Poulvac IB Primer, were administered at the same time to provide protection against Newcastle disease (ND), infectious bursal disease (IBD; Gumboro), and infectious bronchitis (IB), respectively. At week 7 (W7), Poulvac AE was administered for protection against avian encephalomyelitis (AE) and fowl pox (FP). At week 9 (W9), three vaccines, Nobilis ND Clone 30, Nobilis IB 4-91, and Nobilis IB Ma5, were administered at the same time for protection against Newcastle disease and infectious bronchitis. At week 12 (W12), Thymovac was administered for protection against chicken infectious anaemia (CIA) (Figure 1).

**Figure 1.**
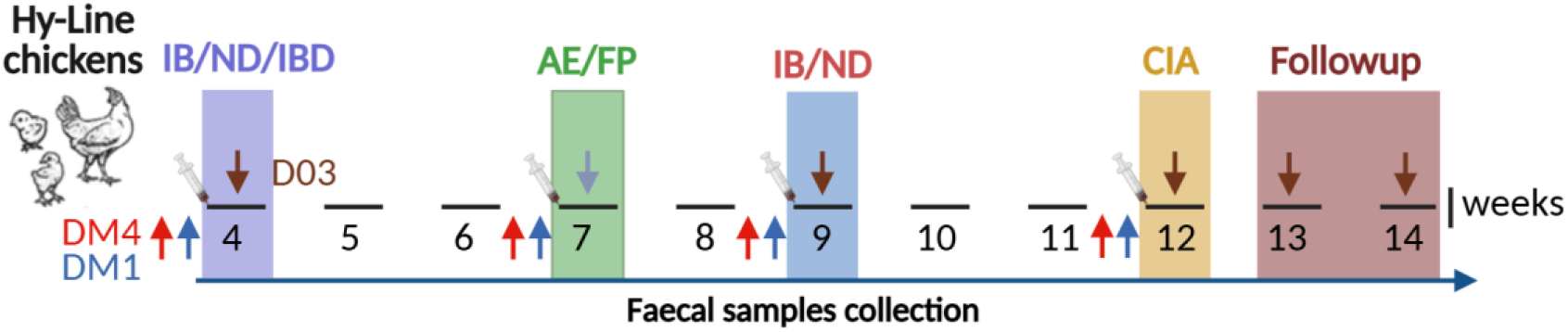
Study design and time points for sample collection from vaccinated Hy-Line chickens. Pooled faecal samples from 3 birds per pen (4 pens in total) were collected longitudinally for different vaccines at three different timepoints: DM4: four days before vaccination; DM1: one day before vaccination; D03: three days after vaccination. Red and blue arrows represent sample collection at 4 days and 1 day before vaccination, respectively, while the brown arrow indicates sample collection at 3 days post-vaccination. The coloured rectangles indicate the vaccination week and the diseases targeted by each vaccine. IB/ND/IBD: infectious bronchitis/Newcastle disease/gumboro was administered at week 4; AE/FP: avian encephalomyelitis/fowl pox was administered at week 7; IB/ND: infectious bronchitis/Newcastle disease was administered at week 9; CIA: chick anaemia was the last vaccine administered at week 12. The followup period at weeks 13 and 14 is defined as a time with no vaccine administration.

For sample collection, birds from the same pen were placed in a clean, sterile bird crate and fresh faecal samples were collected from the crate floor for each bird. The white uric acid layer was carefully avoided while collecting faecal samples to prevent interference during DNA isolation. The three faecal samples collected per pen were pooled by mixing approximately equal volumes from the birds to ensure representative group sampling without precise weighing. Pooled samples were immediately placed on dry ice and subsequently stored at -80 °C until further processing. All faecal samples were collected at the same time of day throughout the study to minimise potential variation in faecal microbiota composition associated with sampling time.

The same process was performed for each pen and during each vaccination week at three different time points: 1) four days before vaccination (DM4); 2) one day before vaccination (DM1); and 3) three days after vaccination (D03), at different time intervals from week 4 to week 12. Faecal samples were also collected at weeks 13 (W13, F1) and 14 (W14, F2) when no vaccine was administered, to determine the temporal variations in the gut microbiota in the following weeks as follow-up. The comparison between time points 1 and 2 provided a short-term pre-vaccination comparison to assess temporal variation in the microbiota before each vaccination. Comparisons between time points 2 and 3 were used to assess changes in the microbiota occurring around the vaccination. Because birds had previously received vaccinations, these time points were not considered an untreated control. Rather, the longitudinal sampling design allowed each pen to serve as its own control, while potential residual effects of previous vaccinations could not be excluded.

### DNA extraction and sequencing

Total DNA from pooled faecal samples (n=56) was isolated using QIAamp PowerFecal Pro DNA Kit (QIAGEN®) following the manufacturer’s instructions. As the chicken faecal samples contain uric acid, which affects DNA quality, an additional DNA purification step was performed using AMPure XP Beads (Beckman Coulter) at a modified 1:1 volume ratio, followed by the remaining steps as per the manufacturer’s instructions. The DNA quality was assessed by 1.2% agarose gel electrophoresis, while DNA quantification was performed using a Nanodrop™ 1000 Spectrophotometer (ThermoFisher Scientific, UK) and Qubit® 4.0 (Invitrogen Life Technologies, Carlsbad, CA, USA). Purified DNA samples were sent to Novogene, UK, for sequencing of the V4 region of the 16S rRNA gene using Universal primers 515F (5′-GTGCCAGCMGCCGCGGTAA-3′) and 806R (5′-GGACTACHVGGGTWTCTAAT-3′). This process generated 250-bp raw reads using the Illumina NovaSeq 6000 platform.

### Bioinformatics analysis

The raw paired-end sequence reads, with primers and adapters removed by Novogene, were quality-checked using FastQC (v0.11.9) (Simon Andrews *et al*. 2010). The reads were then demultiplexed and denoised using the DADA2 plugin implemented in QIIME2 (release 2025.7.0; Quantitative Insights Into Microbial Ecology), generating representative sequences and a feature table. Taxonomic classification was conducted using the SILVA 138.2 reference database (Quast et al., 2013). As no pre-trained classifier was available for this version, a Naive Bayes classifier was trained following the RESCRIPt-based workflow described by Robeson et al. (2021) (Robeson *et al*. 2021). The SILVA 138.2 SSU_NR99 dataset was curated to remove redundant sequences at 99% identity, converted to DNA format, and trimmed to the V4 region corresponding to the 515F/806R primer pair before classifier training. The trained SILVA 138.2 classifier was then used to assign taxonomy to the representative amplicon sequence variant (ASVs). A phylogenetic tree of the representative ASVs was constructed to support diversity analyses. For this, the representative ASV sequences were aligned using MAFFT (Katoh and Standley 2013), poorly aligned positions were masked, and a maximum-likelihood tree was inferred using FastTree (Price, Dehal, and Arkin 2009). The resulting tree was midpoint-rooted for downstream phylogenetic analyses.

Any reads belonging to “Unassigned”, “Archaea”, and “chloroplasts” were removed. The resulting reads were used to create a phyloseq object in RStudio (v4.5.0), and the commonly found environmental contaminants reported to be present in DNA extraction reagents, such as *Nitrospira, Georgenia, Denitratisoma, Luimnobacter*, and *Nitrosomonas* were removed. Sequencing depth was assessed across samples, and rarefaction curves were examined to evaluate the relationship between sequencing depth and observed ASVs. Observed, Shannon and InvSimpson indices were calculated using the estimate_richness function of phyloseq (v1.52.0). An unweighted UniFrac distance-based principal coordinates analysis (PCoA) plot was generated to examine clustering among groups. Bar graphs displaying the top 10 phyla and genera were plotted using ggplot2 (v3.5.2).

### Statistical analysis

Normality of data was assessed using the Shapiro-Wilk test. Alpha diversity indices (Shannon and InvSimpson) were analysed using generalised linear mixed models (GLMMs) implemented in the glmmTMB package. Vaccine, timepoint, and their interaction were included as fixed effects, and pen was included as a random effect to account for repeated measures. Depending on data distribution, a Gaussian family was applied to normally distributed data, whereas a Gamma distribution with a log link was used for non-normally distributed data. Post hoc pairwise comparisons of estimated marginal means were conducted using the emmeans package, with Tukey-adjusted p-values to account for multiple comparisons.

Beta diversity was assessed using unweighted UniFrac distance matrices. Differences in microbial community composition across vaccine groups and timepoints were evaluated using permutational multivariate analysis of variance (PERMANOVA), and visualised using principal coordinates analysis (PCoA).

For differential abundance analysis, GLMMs were fitted for individual taxa to evaluate the effects of vaccine, timepoint, and their interaction, with pen included as a random effect. Only taxa that appear in more than 10% of samples and were detected in more than one vaccine group were retained for analysis. Relative abundance data were log-transformed following the addition of a pseudocount (1×10^−5^) to zero values to stabilise variance (Nishijima *et al*. 2025). Post hoc comparisons were performed using emmeans (Lenth V. Russell and Piaskowski Julia 2025), with Tukey-adjusted pairwise p-values. To account for multiple testing across taxa, Benjamini-Hochberg false-discovery-rate (FDR) correction was additionally applied to the pairwise comparisons within each vaccine group. Only taxa showing non-significant differences in DM4 vs DM1, but significant differences between DM1 and D03, were considered potentially vaccine-associated taxa. This criterion was used to identify taxa that were relatively stable before vaccination but subsequently changed following vaccination. Microbiota stability was further assessed by comparing the final vaccination timepoint (D03) with follow-up timepoints (weeks 13 and 14) using the same modelling framework. P-value ≤ 0.05 was declared significant.

## Results

### Gut microbial diversity exhibited vaccine-dependent temporal shifts

A total of 342,0118 raw paired-end reads were obtained from 56 samples, with an average of 61,703 ± 15,575 reads per sample. After filtering, denoising, merging, and chimera removal, an average of 92.1% of reads were retained, indicating high sequencing quality. The filtered reads were used to generate 2191 unique ASVs. After removing unassigned features and known environmental contaminants, 1866 ASVs were retained for downstream analyses. Observed ASV rarefaction curves approached a plateau, suggesting that sequencing depth was sufficient to capture most of the detectable ASVs in the samples.

To explore how the administration of different vaccines influences gut microbial diversity over time, we compared within-sample (alpha) and between-sample (beta) diversity across vaccine groups and timepoints. The highest alpha diversity was observed in the CIA vaccine group at the DM4 (Shannon: 3.92 ± 0.96; InvSimpson: = 27.7 ± 22.3) timepoint, and the lowest diversity was recorded in the IB/ND/IBD vaccine group at DM1 (Shannon: 1.59 ± 0.22; InvSimpson: = 3.4 ± 0.7) timepoint (Figure 2 A-B). Both indices showed significant effects of vaccines group (Observed: χ^2^ = 48.77, p < 0.001; Shannon: χ^2^ = 35.75, p < 0.001; InvSimpson: χ^2^ = 21.76, p < 0.001), sampling timepoints (Observed: χ^2^ = 9.13, p 0.01; Shannon: χ^2^ = 12.49, p = 0.002; InvSimpson: χ^2^ = 13.19, p = 0.001), and vaccine × timepoints interactions (Observed: χ^2^ = 21.99, p < 0.001; Shannon: χ^2^ = 13.26, p = 0.039; InvSimpson: χ^2^ = 17.76, p = 0.007), indicating temporal vaccine-dependent changes in microbial diversity.

**Figure 2.**
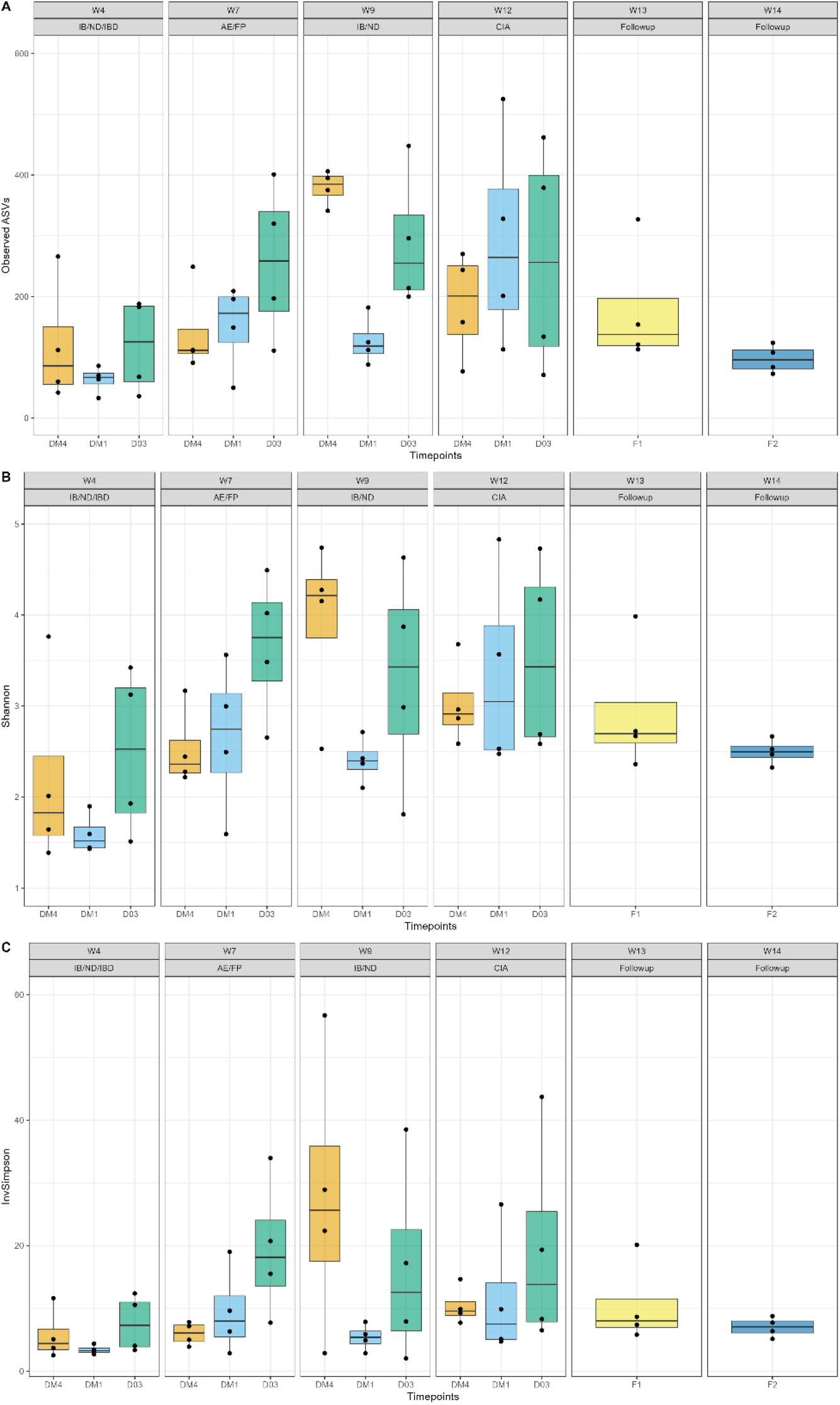
Observed (A), Shannon (B) and InvSimpson (C) diversity indices across different sampling timepoints for each vaccine group. DM4: four days before vaccination; DM1: one day before vaccination; D03: three days after vaccination; F1: follow-up 1 at week 13 (W13); F2: follow-up 2 at week 14 (W14); IB/ND/IBD: infectious bronchitis/Newcastle disease/gumboro at Week 4 (W4); AE/FP: avian encephalomyelitis plus at week 7 (W7); IB/ND: infectious bronchitis/Newcastle disease at week 9 (W9); CIA: chick anaemia at week 12 (W12).

Pairwise comparisons indicated that microbial diversity remained stable during the pre-vaccination period (DM4 vs. DM1) for all vaccine groups except CIA, which showed a significant decrease in Observed ASVs (z = 4.31, p = <0.0001), Shannon (z = 3.278, p = 0.003) and InvSimpson diversity (z = 4.260, p < 0.001). Diversity between the pre-vaccination (DM4) and post-vaccination timepoint (D03) was also stable across all groups, except AE/FP, where both Shannon (Estimated Marginal Mean [EMM], z = -2.360, p = 0.040) and InvSimpson indices (EMM, z = -2.745, p = 0.020) showed a significant increase. Significant decreases in Shannon and InvSimpson diversity were observed between DM1 and D03 timepoints in IB/ND/IBD (EMM, z = -2.64, p = 0.02) and CIA (EMM, z = -2.39, p = 0.04) groups, respectively, whereas AE/FP and IB/ND groups showed no significant changes. A significant decrease in the number of Observed ASVs was also observed in the CIA group (EMM, z = −2.97, p = 0.01).

During the follow-up period after the final CIA vaccination, both Shannon (EMM: D03 vs F1, z = 0.76, p = 0.72; D03 vs F2, z = 1.76, p = 0.18) and InvSimpson (EMM: D03 vs F1, z = 1.01, p = 0.56; D03 vs F2, z = 1.93, p = 0.13) diversity indices showed a non-significant gradual decline. Furthermore, no significant differences were observed between the two follow-up timepoints (F1 vs F2; Shannon: z = 0.57, p = 0.57; InvSimpson: z = 0.92, p = 0.63), supporting the stability of microbial diversity during the post-vaccination period.

To assess changes in overall microbial community structure, beta diversity was evaluated using unweighted UniFrac distances. PERMANOVA analysis revealed a significant effect of vaccine on gut microbial community composition (R^2^ = 0.220, p = 0.001). The effect of sampling timepoints was not significant (R^2^ = 0.039, p = 0.171). However, a significant vaccine × sampling timepoints interaction was observed (R^2^ = 0.138, p = 0.011), indicating that temporal changes in community composition differed among vaccine groups. Ordination analysis using PCoA showed distinct clustering of all the timepoint samples from the IB/ND/IBD group, while no clear separation was observed among the other vaccines or timepoints (Figure 3). Moreover, no significant differences were detected between the final CIA vaccination timepoint (D03) and follow-up timepoints (p ≥ 0.05), suggesting a stabilisation of microbial community structure. Overall, these findings indicate that certain vaccines transiently altered the gut microbiota, but the community composition recovered and stabilised in the subsequent weeks.

**Figure 3.**
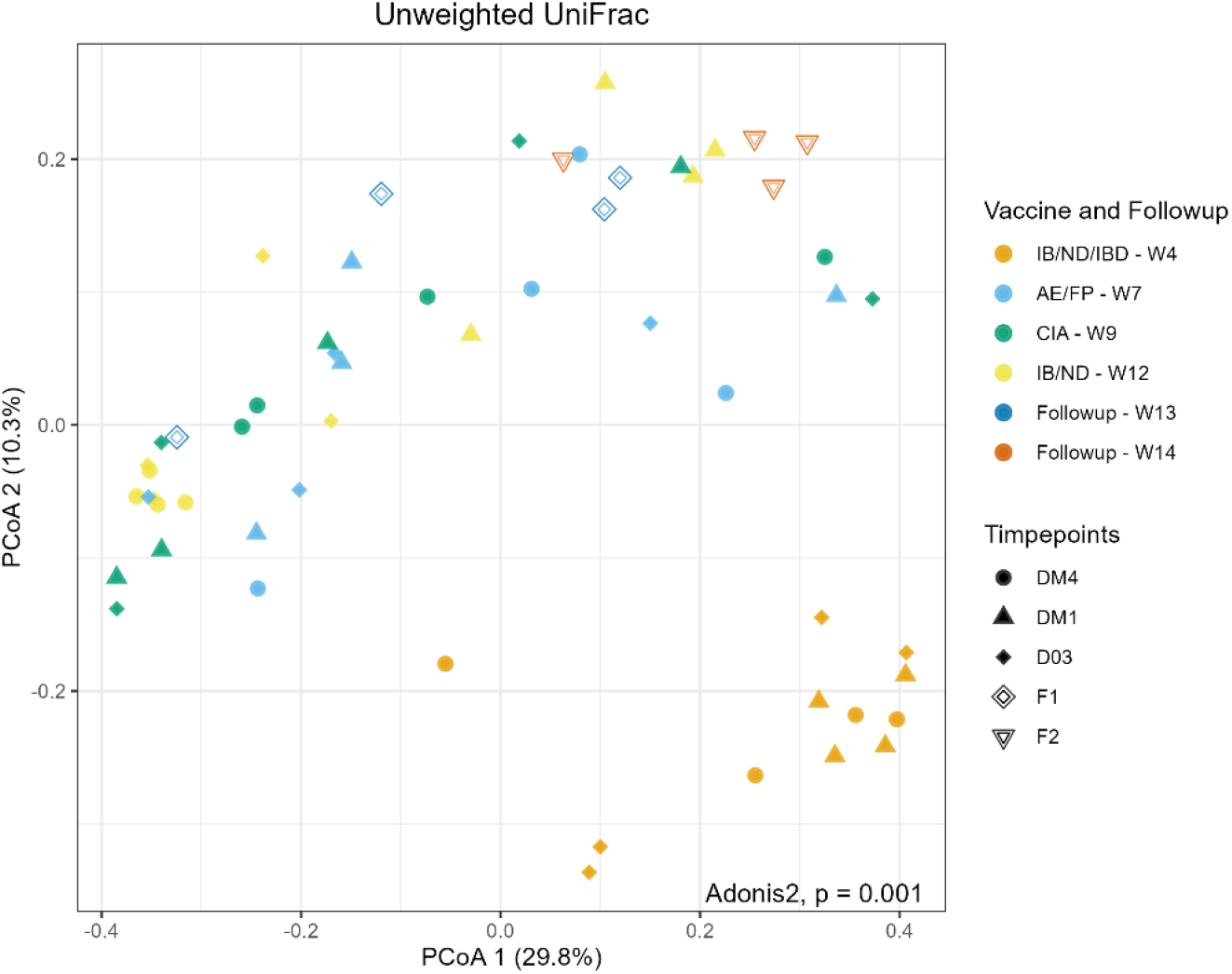
Principal coordinate analysis (PCoA) based on unweighted Unifrac distance displaying clustering of samples across different sampling timepoints for each vaccine group. PCoA 1 indicates the 1st principal component, and PCoA 2 displays the 2nd principal component, altogether showing the total percentage of variation among samples. DM4: four days before vaccination; DM1: one day before vaccination; D03: three days after vaccination; F1: follow-up 1 at week 13 (W13); F2: follow-up 2 at week 14 (W14); IB/ND/IBD: infectious bronchitis/Newcastle disease/gumboro at Week 4 (W4); AE/FP: avian encephalomyelitis plus at week 7 (W7); IB/ND: infectious bronchitis/Newcastle disease at week 9 (W9); CIA: chick anaemia at week 12 (W12).

### Bacillota remained the dominant phylum, while dominant genera varied among vaccine groups

To determine which bacterial taxa were most abundant and how their relative abundances varied across vaccine groups, we analysed gut microbial composition at the phylum and genus levels. We identified 24 phyla and 356 genera across all samples. At the phylum level, Bacillota (65.8 ± 0.10% relative abundance), Fusobacteriota (10.3 ± 0.15%), Bacteroidota (8.85 ± 0.14%), and Pseudomonadota (7.85 ± 0.10%) were highly abundant across all vaccines and timepoints (Figure 4A). Other minor phyla included Actinomycetota (4.63 ± 0.10%) and Verrucomicrobiota (1.50 ± 0.04%). Bacillota remained the dominant phylum across all vaccine groups and during the follow-up weeks, indicating its overall prevalence in the gut microbiota of these chickens.

**Figure 4.**
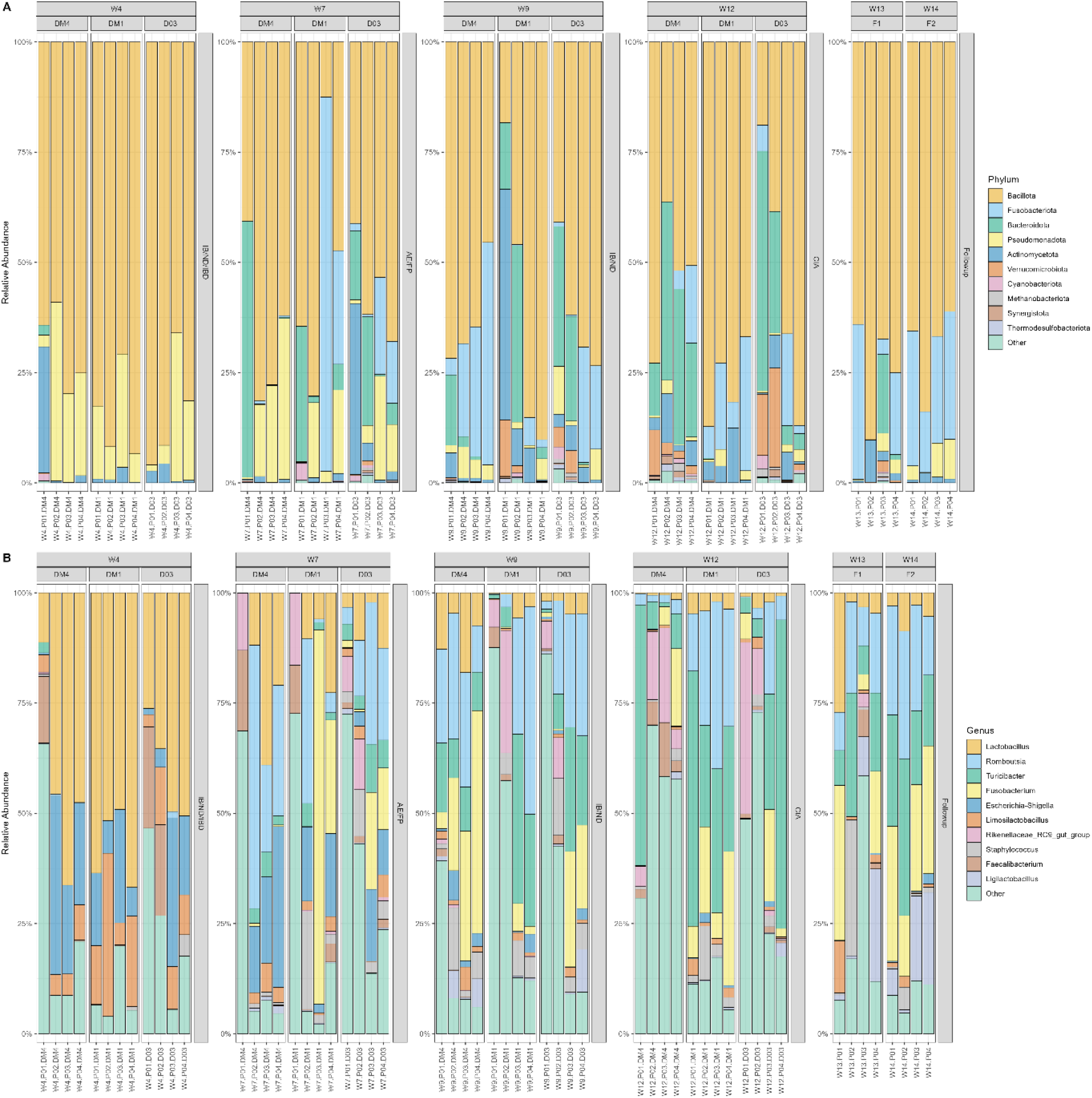
The relative abundance of (A) phyla and (B) genus across different sampling timepoints for each vaccine group. DM4: four days before vaccination; DM1: one day before vaccination; D03: three days after vaccination; F1: follow-up 1 at week 13 (W13); F2: follow-up 2 at week 14 (W14); IB/ND/IBD: infectious bronchitis/Newcastle disease/gumboro at Week 4 (W4); AE/FP: avian encephalomyelitis plus at week 7 (W7); IB/ND: infectious bronchitis/Newcastle disease at week 9 (W9); CIA: chick anaemia at week 12 (W12).

At the genus level, *Lactobacillus* (15.2 ± 0.19%), *Romboutsia* (13.4 ± 0.14%), *Turicibacter* (12.3 ± 0.16%), and *Fusobacterium* (10.3 ± 0.16%) were most abundant (Figure 4B). Other minor genera were *Escherichia-Shigella* (6.52 ± 0.10%), *Limosilactobacillus* (3.76 ± 0.10%), *Rikenellaceae_RC9_gut_group* (3.62 ± 0.10%), and *Staphylococcus* (3.29 ± 0.10%). Approximately 3.87 ± 0.10% ASVs could not be assigned to any known genus and were classified as *Incertae_Sedis*. The dominant genera varied among vaccine groups. *Lactobacillus* predominated in the IB/ND/IBD group, whereas *Romboutsia* was more abundant in the AE/FP group. In the IB/ND group, *Romboutsia* and *Turicibacter* were dominant, and in the CIA group, *Turicibacter* became the most abundant genus. During the follow-up weeks, *Fusobacterium* also became abundant, alongside *Romboutsia* and *Turicibacter*.

### Vaccine-associated variation was observed at the genus level despite phylum-level stability

To investigate vaccine-specific temporal effects on gut microbiota, we examined differentially abundant taxa within each vaccine group, focusing on microbial taxa that significantly changed post-vaccination (DM1 vs D03), remained stable during the pre-vaccination baseline (DM4 vs DM1) to capture a potential vaccine effect (Figure 5).

**Figure 5.**
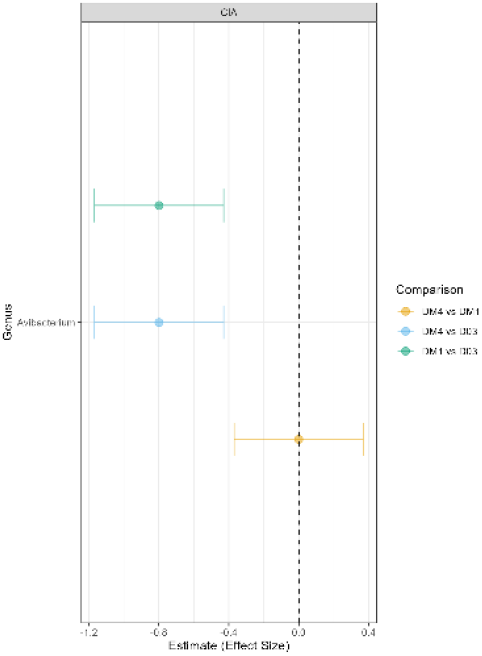
Forest plot showing the estimated effect size of differentially abundant genera in the CIA vaccine group. Points represent the estimated effect size of each genus, with 95% confidence intervals shown as horizontal error bars. Values to the left of the vertical line indicate decreases in the relative abundance, while values to the right show increases in the relative abundance. DM4: four days before vaccination; DM1: one day before vaccination; D03: three days after vaccination; CIA: chick anaemia vaccine administered at week 12.

Using these criteria, no phyla were identified as differentially abundant within each vaccine group. At the genus level, *Avibacterium* (EMM, t = -4.24, p = 0.01) showed a significant decrease in the CIA group. No significantly different genera were identified in other vaccine groups.

During follow-up (weeks 13 and 14), no significantly differentially abundant taxa (p > 0.05) were observed compared with the post-vaccination timepoint (D03), suggesting overall stability of the gut microbial community.

## Discussion

Vaccines are recognised as an important factor influencing the gut microbiota of chickens (Redweik *et al*. 2020). In commercial production systems, birds are routinely exposed to multiple vaccines through drinking water to protect them against a broad range of pathogens. Oral vaccines potentially engage with the gut mucosal immune system by delivering antigens to the specialised intestinal lining while interacting closely with the gut microbial environment (Wang *et al*. 2026). In this longitudinal study, we examined gut microbial dynamics in response to a series of live and inactivated vaccines administered at different timepoints from early life through sexual maturity, mimicking a commercial vaccine regimen for layer production. To our knowledge, this is the first longitudinal study to investigate microbiome responses to complex, multi-vaccine programs mimicking commercial conditions.

In this study, microbial diversity remained relatively stable across most vaccine groups, indicating a notable resilience of the layer chicken gut microbiota despite multiple immunological interventions. Previous studies have reported variable effects of vaccination on gut microbiota. Some studies observed an increase in diversity post-vaccination, whereas others reported minimal influence, highlighting the complexity of host-microbe-vaccine interactions (Beck, Zhao, and Erf 2024). In contrast to the overall stability, we observed a decrease in alpha diversity in IB/ND/IBD and CIA groups post-vaccination. This finding aligns with a recent report showing that IBV vaccination was associated with reduced bacterial diversity (Borey *et al*. 2021). The CIA vaccine in this study is a live attenuated formulation. Live vaccines have been reported to induce greater changes in gut microbial composition than inactivated vaccines, potentially through transient colonisation, local immune activation, or niche competition (Loddo *et al*. 2025). However, live attenuated *Salmonella Typhimurium* vaccination in layer chickens exerted minimal effect on alpha diversity, likely reflecting differences in vaccine type, target pathogen, or host response (Khan *et al*. 2024).

Interestingly, we also observed fluctuations in microbial diversity during the pre-vaccination period in some vaccine groups. These shifts likely reflect natural microbiome succession associated with birds’ age and environmental influences, such as diet, stress, and housing, consistent with previous findings (Wickramasuriya *et al*. 2022). Importantly, because of pre-vaccination temporal variation, we cannot exclude the possibility that part of the post-vaccination variation reflects ongoing natural fluctuations rather than vaccine effects alone. However, our statistical framework accounted for both sampling timepoints and vaccine × timepoints interactions, allowing us to distinguish vaccine-associated deviations from shared temporal trends. Such baseline dynamics highlight the importance of including pre-vaccination sampling in microbiome studies. Without accounting for these natural successional changes, observed post-vaccination differences may reflect a combination of vaccination effects and underlying temporal variation, making it difficult to attribute changes solely to vaccination. Importantly, after the final vaccine dose, microbial diversity gradually recovered by weeks 13-14, with no significant further change between follow-up timepoints, indicating that the microbiome largely stabilised post-vaccination.

At the taxonomic level, phyla Bacillota and Fusobacteriota were dominant across all vaccine groups and sampling timepoints, consistent with the typical faecal microbiota composition of layer chickens reported previously (Feng *et al*. 2025). We didn’t find any differentially abundant phyla, suggesting that vaccination did not induce major shifts at higher taxonomic levels. At the genus level, we observed different dominant genera during different weeks. In contrast, genus-level profiles revealed marked temporal shifts in dominant taxa. These changes appeared more closely associated with host age than with vaccination, as earlier studies similarly reported high *Lactobacillus* abundance during early life, followed by replacement with *Romboutsia* around week 8 and increasing *Turicibacter* in later stages of growth (Song *et al*. 2024). These age-responsive genera are generally linked to gut development, improved nutrient utilisation, and enhanced immune maturation (Song *et al*. 2024, Liu Q *et al*. 2025).

Differential abundance analysis identified limited evidence of vaccination-associated changes in the gut microbiota. No phyla showed differential abundance within any vaccine group. At the genus level, *Avibacterium* showed a decrease following vaccination in the CIA group, while no genera met the predefined criteria in the other vaccine groups. *Avibacterium* is a member of the Pasteurellaceae family and has been reported as a member of the avian respiratory and gastrointestinal microbiota (Blackall 2020). However, the biological significance of the observed change in relative abundance is unclear, particularly given that this was a pilot study with a limited number of pen-level samples. Furthermore, the exploratory nature of the analysis and the lack of consistent differential abundance across vaccine groups mean that this finding should not be interpreted as definitive evidence of a vaccine-driven microbial shift.

A limitation of this study is the use of faecal samples rather than caecal content samples for gut microbiota analysis. While faecal microbiota profiling provides valuable insights into intestinal communities, it may not fully capture mucosa-associated microbial populations and can be influenced by transient environmental or sampling variability. Moreover, faecal samples are not fully representative of the caecal microbiota, where the highest microbial density and diversity in chickens is typically observed (Rychlik 2026). However, obtaining caecal contents from live birds is not feasible in a longitudinal experimental design; therefore, faecal sampling was used to enable repeated monitoring of microbiome dynamics over time. Consequently, some potential vaccine-associated microbial changes may be attenuated or not fully reflected in faecal microbial profiles. Another limitation of this study was the relatively small number of biological replicates, with only four pen-level samples included in the analysis. This limited sample size may have reduced statistical power, increasing the likelihood that biologically relevant differences in microbial composition or diversity between groups and over time were not detected. In this study, the findings were considered exploratory rather than definitive evidence of a vaccine effect. Future studies incorporating a larger number of replicates from individual birds would be helpful in capturing the vaccine-associated changes with improved statistical power and greater confidence.

## Conclusion

This longitudinal study investigated the impact of various vaccines administered to layer chickens at different ages within a commercial setting on the gut microbial community. Our findings revealed that microbial diversity remained relatively stable across most vaccine groups. Differential abundance analyses didn’t identify significant changes in taxa following vaccination. Moreover, the microbial community showed stability following the vaccination period. This work offers a foundational framework for future studies on microbiome-vaccine interactions and their implications for poultry health, immune development, and productivity.

## Data availability statement

The datasets generated during the current study are available in the European Nucleotide Archive under the project number PRJEB104076 (https://www.ebi.ac.uk/ena/browser/view/PRJEB104076).

## Funding information

This work was supported by a BBSRC Institute Development Grant through ECR pump priming 24/25 grant funding (supported through the Institute Strategic Program BBS/E/RL/230001A) secured by Anum Ali Ahmad. Laura Glendinning is supported by a University of Edinburgh Chancellor’s Fellowship. For the purpose of open access, the author has applied a Creative Commons Attribution (CC BY) licence to any Author Accepted Manuscript version arising from this submission.

## Acknowledgements

We would like to thank the staff at the National Avian Research Facility for their help in the management of birds. We gratefully acknowledge strategic investment by the Biotechnology & Biological Sciences Research Council to The Roslin Institute for 2023-28.

## Contributions

Anum Ali Ahmad (Conceptualisation, Methodology, Funding acquisition, Investigation, Supervision, Project administration, Formal analysis, Writing – Original Draft, Writing – Review & Editing), Kris Hogan (Resources, Writing - Review & Editing), Laura Glendinning (Methodology, Writing – Review & Editing).

## Conflicts of interest

All authors affirm that they have no conflicts of interest to declare.

## Notes

https://www.ebi.ac.uk/ena/browser/view/PRJEB104076

### Competing Interest Statement

The authors have declared no competing interest.

### Summary of Updates

Figures 2, 3, 4, and 5 are revised along with the methodology.

## References

Akhtar MS, Zaman W. From gut to shot: Microbiome-guided strategies to improve vaccine responses in food animals. Vaccines (Basel) 2026;14(4):327. 10.3390/vaccines14040327.

Beck CN, Zhao J, Erf GF. Vaccine immunogenicity versus gastrointestinal microbiome status: Implications for poultry production. Applied Sciences 2024;14(3):1240. 10.3390/app14031240.

Blackall PJ. Avibacterium. In: Christensen H (ed.), Bergey’s Manual of Systematics of Archaea and Bacteria. n.p.: Wiley, 2020, 1–9. 10.1002/9781118960608.gbm01840.

Borey M, Bed’Hom B, Bruneau N et al. Caecal microbiota composition of experimental laying hens differs according to genetic line and vaccination to IBV. Preprint, 9 Feb. 2021. 10.21203/rs.3.rs-177864/v1.

Das Q, Shay J, Gauthier M et al. Effects of vaccination against Coccidiosis on gut microbiota and immunity in Broiler fed bacitracin and berry pomace. Front Immunol 2021;12. 10.3389/fimmu.2021.621803.

FAO. Food Outlook – Biannual Report on Global Food Markets. Rome, 2025. 10.4060/cd7448en.

Feng Z, Lorenc N, O’Brien B et al. Deep culturing the fecal microbiota of healthy laying hens. Anim Microbiome 2025;7(1):32. 10.1186/s42523-025-00395-y.

Katoh K, Standley DM. MAFFT multiple sequence alignment software version 7: Improvements in performance and usability. Mol Biol Evol 2013;30(4):772–80. 10.1093/molbev/mst010.

Khan S, McWhorter AR, Andrews DM et al. A live attenuated Salmonella Typhimurium vaccine dose and diluent have minimal effects on the caecal microbiota of layer chickens. Front Vet Sci 2024;11. 10.3389/fvets.2024.1364731.

Khan S, McWhorter AR, Willson NL et al. Vaccine protection of broilers against various doses of wild-type Salmonella Typhimurium and changes in gut microbiota. Veterinary Quarterly 2025;45(1):1–14. 10.1080/01652176.2024.2440428.

Lenth V. Russell and Piaskowski Julia. Emmeans: Estimated Marginal Means, Aka Least-Squares Means. 2025. https://CRAN.R-project.org/package=emmeans (10 Nov. 2025, date last accessed).

Liu PY, Liaw J, Soutter F et al. Multi-omics analysis reveals regime shifts in the gastrointestinal ecosystem in chickens following anticoccidial vaccination and Eimeria tenella challenge. MSystems 2024;9(10). 10.1128/msystems.00947-24.

Liu Q, Akhtar M, Kong N et al. Early fecal microbiota transplantation continuously improves chicken growth performance by inhibiting age-related Lactobacillus decline in jejunum. Microbiome 2025;13(1):49. 10.1186/s40168-024-02021-6.

Loddo F, Laganà P, Rizzo CE et al. Intestinal microbiota and vaccinations: A systematic review of the literature. Vaccines (Basel) 2025;13(3):306. 10.3390/vaccines13030306.

Nishijima S, Stankevic E, Aasmets O et al. Fecal microbial load is a major determinant of gut microbiome variation and a confounder for disease associations. Cell 2025;188(1):222–236.e15. 10.1016/j.cell.2024.10.022.

Price MN, Dehal PS, Arkin AP. FastTree: Computing Large Minimum Evolution Trees with Profiles instead of a Distance Matrix. Mol Biol Evol 2009;26(7):1641–50. 10.1093/molbev/msp077.

Redweik GAJ, Daniels K, Severin AJ et al. Oral treatments with probiotics and live Salmonella vaccine induce unique changes in gut neurochemicals and microbiome in chickens. Front Microbiol 2020;10. 10.3389/fmicb.2019.03064.

Robeson MS, O’Rourke DR, Kaehler BD et al. RESCRIPt: Reproducible sequence taxonomy reference database management. PLoS Comput Biol 2021;17(11):e1009581. 10.1371/journal.pcbi.1009581.

Rychlik I. Threats and Opportunities When Using Chickens as a Model for Host–Microbiota Studies. Microorganisms 2026;14(6):1330. 10.3390/microorganisms14061330.

Shterzer N, Rothschild N, Sbehat Y et al. Vertical transmission of gut bacteria in commercial chickens is limited. Anim Microbiome 2023;5(1):50. 10.1186/s42523-023-00272-6.

Simon Andrews FK, Segonds-Pichon A, Biggins L et al. FastQC: a quality control tool for high throughput sequence data. Preprint, 2010. http://www.bioinformatics.babraham.ac.uk/projects/fastqc (10 Nov. 2025, date last accessed).

Simon K, Verwoolde MB, Zhang J et al. Long-term effects of early life microbiota disturbance on adaptive immunity in laying hens. Poult Sci 2016;95(7):1543–54. 10.3382/ps/pew088.

Song B, Sun P, Kong L et al. The improvement of immunity and activation of TLR2/NF-κB signaling pathway by Romboutsia ilealis in broilers. J Anim Sci 2024;102. 10.1093/jas/skae286.

Wang X, Pan W, Huang J et al. Oral vaccine delivery systems: Gastrointestinal barriers, gut microbiota, and emerging engineering strategies. Cell Rep 2026;45(10):118038. 10.1016/j.celrep.2026.118038.

Wickramasuriya SS, Park I, Lee K et al. Role of physiology, immunity, microbiota, and infectious diseases in the gut health of poultry. Vaccines (Basel) 2022;10(2):172. 10.3390/vaccines10020172.

Wiertsema SP, Bergenhenegouwen J van, Garssen J et al. The interplay between the Gut Microbiome and the Immune System in the Context of Infectious Diseases throughout Life and the Role of Nutrition in Optimizing Treatment Strategies. Nutrients 2021;13(3):886. 10.3390/nu13030886.

